# Regulators of proteostasis are translationally repressed in fibroblasts from sporadic and LRRK2-G2019S Parkinson’s patients

**DOI:** 10.1101/2022.01.17.476095

**Authors:** Dani Flinkman, Ye Hong, Prasannakumar Deshpande, Sirkku Peltonen, Valtteri Kaasinen, Peter James, Eleanor T. Coffey

## Abstract

Parkinson’s disease (PD) is characterized by heterogeneous motor and non-motor symptoms. The majority of PD cases are sporadic, whereas the leucine-rich repeat kinase 2 (LRRK2) G2019S mutation is the most common monogenic cause of familial PD and symptomatically indistinguishable from the sporadic form. Deficits in protein synthesis have been reported in PD. However, it is not known which proteins are affected or if there are synthesis differences between patients with sporadic and LRRK2 G2019S PD. Here we used non-canonical amino acid labeling and mass-spectrometry based proteomics to perform a quantitative comparison of nascent proteins in fibroblasts from sporadic and G2019S mutation PD patients compared to healthy individuals. We found that several proteins were under-represented among nascent proteins in cells from PD patients. Among these were regulators endo-lysosomal sorting, mRNA processing and translation itself. We further validated which of these proteins were dysregulated at the static proteome level using targeted proteomics. Key downregulated proteins included the autophagy regulator ATG9A and translational stability regulator YTHDF3. Notably, 68% of the identified proteins were downregulated in both sporadic and LRRK2-G2019S forms of PD. We identify translational repression of key regulators of proteostasis in patient fibroblasts that are common to both sporadic and LRRK2-G2019S PD.

## INTRODUCTION

Parkinson’s disease (PD) is the fastest growing neurodegenerative disease, with an estimated 6.1 million patients worldwide in 2016 (Collaborators, 2018). The cardinal motor symptoms include bradykinesia, muscular rigidity, resting tremor and postural instability, but the disease is often complicated by non-motor symptoms, such as hyposomia, constipation and REM sleep behavior disorder, which can precede disease onset by years (Kalia and Lang, 2015). However, the disease is heterogenous, and phenotypes of motor and non-motor symptoms together with general disease trajectories differ broadly among patients (Berg, et al., 2021). Moreover, the diagnosis itself is error prone as the core motor symptoms can be difficult to distinguish from those of other movement disorders (Titova, et al., 2017). There is a growing interest in exploiting ‘omic’ technologies to understand the molecular changes that contribute to the pathology and its heterogeneity, and to identify markers that could potentially assist in the diagnosis and stratification of PD patients, ideally at an early stage (Parnetti, et al., 2019,Schilder, et al., 2021).

In recent years, studies have identified disease-associated features of PD in patient skin fibroblasts. For instance, alpha-synuclein seeding activity measured from PD cadaver skin has been used to successfully distinguish patients with synucleinopathies (Doppler, 2021,Orrù, et al., 2021,Wang, et al., 2020). Also, sebum collected from PD patient’s skin was shown to contain disease defining lipid biomarkers (Sinclair, et al., 2021). We have previously shown that *de novo* protein synthesis is reduced in fibroblasts from PD patients with a receiver operating characteristic (ROC) area under the curve value of 0.925, suggesting good predictive power (Deshpande, et al., 2020). Consistent with this, in a recent study of PD patient-derived organoid cultures of fibroblast origin, a transcriptome analysis identified “cytoplasmic ribosomal proteins” and “translation factors” as the most enriched features (Trapecar, et al., 2021).

In this study, we isolated *de novo* synthesized proteins from patient skin cells to determine if we could identify if particular proteins showed altered translation in Parkinson patient skin cells. We did this for sporadic patient and healthy volunteer samples collected from the Southwest Finland and for LRRK2-G2019S patients, obtained from biorepositories. We used biorthogonal non-canonical amino acid tagging (BONCAT) to label the nascent proteins followed by LC-MS/MS-based analysis (Dieterich, et al., 2006). We found a significant decrease in nascent proteins in both PD groups. Among these, we detected 22 proteins that remained significantly changed (FDR<0.05) at the level of the whole cell proteome. These proteins were not recovered by homeostatic mechanisms and remained substantially lower than in cells from healthy individuals. They represented regulators of various steps of the mRNA translation program and autophagy. These data suggest that deregulated translation can be observed in fibroblasts derived from PD patient skin. Among the proteins that are disturbed at the level of protein synthesis are regulators of endo-lysosomal sorting and of the core translation mechanism itself.

## METHODS

### Cultivation of fibroblasts from patient skin biopsies

Skin punches were taken from the upper arm of donors and patients by a licensed dermatologist at Turku University Hospital (TUH). Punches were placed immediately in Minimal essential medium (M.E.M., Sigma Aldrich). Tissue was chopped into several small pieces using a sterile blade and incubated in M.E.M. supplemented with Gln (2 mM), penicillin (50 U/ml) and streptomycin (50 μg/ml) at 37°C, 5% CO2. Fibroblasts were passaged when 95% confluent. Experiments were done at passage 7 to 9.

### Patient details

The fibroblasts used in this study came from three sources. All sporadic cases were from Turku University Hospital. These included 13 sporadic and 7 age-matched healthy cases from whom skin punches were obtained by a registered dermatologist from the arm. These patients were diagnosed as having PD by neurological testing by a movement disorder specialist using MDS clinical diagnostic criteria for PD combined with dopamine transporter SPECT imaging with [^123^I]FP-CIT as tracer. For LRRK2-G2019S patients, we obtained fibroblasts from the National Institute of Neurological Disorders (NINDS) repository and from the Telethon Network of Genetics Biobanks (TNGB) for 4 LRRK2-G2019S and 14 healthy samples. Patient details including age, sex, motor MDS-UPDRS, and Hoehn & Yahr stage, as well as [^123^I]FP-CIT SPECT data for striatal dopamine transporter binding are described in Table 1. All patient samples were obtained with appropriate permissions and informed consents in accordance with the ethical guidelines from the Turku University Hospital and from the NINDS repository (USA).

**Table 1.** Demographic and clinical characteristics of patients and healthy controls. Fibroblasts from Parkinson’s disease (PD1-PD13) patients with sporadic disease, or healthy donors (C1-C7) from Turku University Hospital (T.U.H.) in Finland are shown alongside LRRK2-G2019S patient and healthy subject fibroblasts from the National Institute of Neurological Disorders (NINDS) and the Biobank (EMB) are shown. Gender, male (m) or female (f) is indicated and age at the time of biopsy are shown. *Unified Parkinson’s Disease rating scale (UPDRS).

| Cell I.D. | Description | Source | Gender | Age at sampling | Estimated age of onset | Motor symptom duration (years) | Hoehn & Yahr stage | Motor MDS-UPDRS (Part III)* | DAT binding defect with[123I] FP-CIT |
| --- | --- | --- | --- | --- | --- | --- | --- | --- | --- |
| PD1 | Sporadic | T.U.H. | M | 60 | 56 | 4 | 2 | 50 | Not performed |
| PD2 | Sporadic | T.U.H. | M | 69 | 60 | 9 | 2 | 42 | Not performed |
| PD3 | Sporadic | T.U.H. | M | 59 | 49 | 10 | 3 | 57 | yes |
| PD4 | Sporadic | T.U.H. | M | 52 | 50 | 1.5 | 1 | 25 | Not performed |
| PD5 | Sporadic | T.U.H. | M | 64 | 59 | 5 | 2 | 30 | Not performed |
| PD6 | Sporadic | T.U.H. | M | 65 | 61 | 4 | 2 | 37 | yes |
| PD7 | Sporadic | T.U.H. | M | 69 | 58 | 11 | 2 | 25 | Not performed |
| PD8 | Sporadic | T.U.H. | M | 68 | 65 | 3 | 2 | 32 | yes |
| PD9 | Sporadic | T.U.H. | M | 66 | 64 | 2 | 2 | 32 | yes |
| PD10 | Sporadic | T.U.H. | M | 49 | 46 | 3 | 1 | 26 | yes |
| PD11 | Sporadic | T.U.H. | M | 60 | 50 | 10 | 3 | 47 | yes |
| PD12 | Sporadic | T.U.H. | F | 66 | 56 | 10 | 2 | 15 | yes |
| PD13 | Sporadic | T.U.H. | M | 63 | 48 | 15 | 3 | 20 | Not performed |
| 29492 | LRRK2 G2019S | NINDS | M | 72 | 58 | 14 | 2 | - | - |
| 32949 | LRRK2 G2019S | NINDS | M | 81 | 75 | 6 | 1,5 | 14 | - |
| 32973 | LRRK2 G2019S | NINDS | M | 72 | 62 | 10 | - | - | - |
| 30116 | Sporadic | NINDS | F | 67 | 43 | 24 | - | - | - |
| 30159 | Sporadic | NINDS | F | 76 | 72 | 4 | 2 | - | - |
| 32462 | Sporadic | NINDS | M | 75 | 74 | 1 | 2 | - | - |
| FF0212011 | LRRK2 G2019S | EMB | M | 47 | 38 | 9 | - | - | - |
| FF0432012 | LRRK2 G2019S | EMB | M | 57 | 54 | 3 | - | - | - |
| FF0642009 | LRRK2 G2019S | EMB | F | 58 | 41 | 17 | - | - | - |
| FF0592012 | Sporadic | EMB | F | 69 | 58 | 11 | - | - | - |
| FF0372011 | Sporadic | EMB | M | 40 | 38 | 2 | - | - | - |
| C1 | Healthy subject | T.U.H. | F | 57 | - | - | - | - | - |
| C2 | Healthy subject | T.U.H. | F | 54 | - | - | - | - | - |
| C3 | Healthy subject | T.U.H. | F | 66 | - | - | - | - | - |
| C4 | Healthy subject | T.U.H. | M | 58 | - | - | - | - | - |
| C5 | Healthy subject | T.U.H. | F | 49 | - | - | - | - | - |
| C6 | Healthy subject | T.U.H. | M | 73 | - | - | - | - | - |
| C7 | Healthy subject | T.U.H. | F | 65 | - | - | - | - | - |
| 34770 | Healthy subject | NINDS | M | 72 | - | - | - | - | - |
| 36320 | Healthy subject | NINDS | F | 71 | - | - | - | - | - |
| 35044 | Healthy subject | NINDS | M | 77 | - | - | - | - | - |
| F0172014 | Healthy subject | EMB | F | 68 | - | - | - | - | - |
| F0492012 | Healthy subject | EMB | M | 41 | - | - | - | - | - |
| F0502011 | Healthy subject | EMB | F | 63 | - | - | - | - | - |
| F0682013 | Healthy subject | EMB | F | 58 | - | - | - | - | - |

### AHA labeling of fibroblasts and fluorescence quantitation

AHA labelling was carried out as previously (Deshpande, et al., 2020). Briefly, skin cells were plated at a density of 50,000 cells per well on round coverslips (1.3 cm diameter). At 48 hours post-plating, cells were washed 1× and incubated with Met-free DMEM (Gibco, Thermo Fisher Scientific) supplemented with 2 mM Gln 30 min. To label newly synthesized proteins, we incubated cells for 30 min in the Met-free media containing 1 mM l-azidohomoalanine (AHA). Cells were then washed with 1 ml of PBS and fixed by adding 4% paraformaldehyde. Cells were permeablized overnight in *block* (0.25% Triton X100, 0.2% BSA, 5% sucrose, and 10% horse serum in PBS), and washed 5× with 1 mL of 3% BSA in PBS. We carried out cycloaddition of Alexa-488 to AHA labelled proteins using the Thermo Fisher Scientific kit (catalogue # A10267) according to the manufacturer’s instructions. Nuclei were stained with 1:2000 Hoechst-33342 in PBS. Cells were imaged using a 40× objective and a Leica DMRE microscope with an ORCA C4742-95 CCD camera (Hamamatsu). Images were acquired using identical settings for all samples. Mean fluorescence intensity r.o.i.s in the soma were measured from all cells using ImageJ. The analysis was performed by an experimenter that was blinded to the treatment until after the analysis was completed.

### AHA enrichment and sample preparation for LC-MS/MS

Patient fibroblasts were plated at a density of 400,000 on 10 cm dishes in 10 ml of Dulbeco’s Modified Essential Medium (D.M.E.M.) supplemented with 10% fetal bovine serum (FBS), 2 mM Gln, penicillin (50 U/ml) streptomycin (50 μg/ml) and incubated at 37 °C in 5 % C02 for 6 days. Before AHA labelling, medium was replaced with methionine-free RPMI 1640 medium supplemented with 10 % dialyzed FBS (A338201, ThermoFisher Scientific; filter-sterilized using a 0.45 um filter), 2 mM Gln and incubated for 60 min at 37 °C in 5 % C02. L-azidohomoalanine (AHA) (BCAA005-500, Baseclick) was added to the medium to a final concentration of 4 mM and cells were incubated for a further 18 h. AHA-labelled cells were collected by scraping, and pelleted at 4000 RCF for 5 min at 4 °C. Cells were lysed with lysis buffer (20 mM HEPES pH 7.4, 2 mM EGTA, 1% SDS, 50 mM β-glycerophosphate, 1 mM DTT, 1 mM Na_3_VO_4_, 1% Triton X-100, 10 % Glycerol, 50 mM NaF, 1mM Benzamidine, 1 μg/ml of Aprotonin, Leupeptin and Pepstatin 100 μg/ml PMSF). DNA was digested with the addition of 250 U/ml Benzonase (#70747, Merck, KGaA, Darmstadt, Germany), and by incubation on ice for 15 min. The resulting lysate was pre-cleared by centrifugation 13,400 RCF for 5 min at 4 °C, and the supernatant was kept. Protein concentration was determined using a Pierce 660 nm kit with ionic detergent compatibility reagent (Thermo Fisher Scientific Scientific, Inc., Waltham, USA). Supernatants underwent a cyclo-addition reaction to attach AHA-labelled proteins to alkyne agarose using the Click IT cell reaction buffer kit (#C10269, ThermoFisher Scientific). Alkyne agarose (25 μl per sample) was washed with milli-Q and pre-blocked with 2% PVP-40 (Sigma) in *buffer 1* (100 mM Tris pH 8.5, 1 % SDS, 250 mM NaCl, 5 mM EDTA) and 50 μl of click reaction mix was added to the beads followed by addition of 400 ug of lysate. Samples were incubated for 20 h at room temperature with gentle rotation. Post-click reaction beads were washed with milli-Q and incubated at 70 °C for 15 minutes with 1 ml *buffer 1* supplemented with 1 mM DTT. The beads were washed with *buffer 1* and incubated in dark for 30 minutes with *buffer 1* containing 2 mM iodoacetamide with gentle rotation.

Following alkylation, the resin was washed with 10 ml of milli-Q followed by 2 x 10 ml washes with *buffer 1* for 30 min each. This was followed by 2 x 10 ml milli-Q washes for 2 min. Resin was then washed with *buffer 2* (100 mM Glycine pH 2.5, 1 % SDS). *Buffer 2* was removed by washing 2 x with 10 ml of milli-Q for 2 min wash and stored in *buffer 1* until use. SDS was washed away by 1 x 10 ml of 8 M Urea, 1 x 10 ml 20 % Isopropanol followed by 1 x 10 ml 20 % Acetonitrile (ACN) washes. Resin was transferred to a tube in 1 ml of 50 mM Ammonium bicarbonate containing 10 % ACN, and then resuspended in 200 μl of the same solution. Digestion was started by adding 0.5 μg of sequencing grade modified trypsin (Promega Corporation, Madison, USA), and proceeded for 16 h at 37 °C with mixing. Digest was harvested, and resin washed with 500 μl milli-Q and combined with digest. The digest was then dried down with a SpeedVac (Thermo Fisher Scientific), and filtered through Whatman Grade 3 filter material and cleaned-up with C18-UltraMicroSpin columns (The Nest Group, Inc., Southborough, USA). The peptides were then dried and dissolved in 0.1 % Formic acid (FA). Peptide concentration was measured using a Nanodrop nd-1000 (Thermo Fisher Scientific) at 220 nm, and peptide concentrations for LC-MS/MS loading was normalized.

### Sample preparation for PRM analysis

Fibroblast were cultivated in DMEM as described above and lysed in Laemmli sample buffer. Protein concentration was determined using Pierce 660 nm kit with ionic detergent compatibility reagent (Thermo Fisher Scientific), and 12 μg of each sample was loaded on a 12.5% SDS-PAGE gel and run ~1 cm into the resolving gel. Gels were stained with GelCode Blue and samples were excised and cut in to 1×1 mm pieces. Gel pieces were destained with 50 mM ammonium bicarbonate in 50% ACN and dried completely in Speedvac. Pieces were rehydrated in 100 mM ammonium bicarbonate containing 10 mM DTT and incubated 1 h at 56 °C. 55 mM iodoacetoamide in equal volume of 100 mM ammonium bicarbonate was added to gel pieces and incubated 1 h at RT in dark. Following this, excess volume was removed and gel pieces washed with 100 mM ammonium bicarbonate, followed by 100 % ACN wash, wash sequence was repeated once. Gels were completely dehydrated with 100% ACN and dried in speedvac. Gel pieces were reswelled in 50 mM ammonium bicarbonate containing trypsin on ice for 30 min (final trypsin:sample ratio 1:50) and incubated overnight at 37 °C. Peptides were extracted 2x with 200 μl 50% ACN containing 0.5% FA, and 1x with 100 % ACN. Extracts were pooled and dried completely in the Speedvac and stored at −80°C until analysis. PRM assay design included peptides that were significantly changing in G2019S verses healthy and sporadic verses healthy in the AHA dataset. These were based on a union of significantly changing peptide intensities derived from both raw and LFQ intensity values. Synthetic peptides for the chosen targets were ordered from JPT Peptide Technologies (Berlin, Germany). A spectral library was generated with Mascot (Matrix Science Inc, Boston USA). PRM data was designed and analysed in Skyline 4.2.0.19072.

### LC-MS/MS

For AHA labelled samples, a LTQ-Orbitrap (Thermo Fisher Scientific) connected to an Eksigent nanoLC plus HPLC system (Eksigent technologies, Dublin, USA) was used. Constant flow of 10 μl/min was maintained for peptide loading onto a pre-column (PepMap 100; Thermo Fisher Scientific). Peptides were then separated on a 10 mm PicoTip^™^ fused silica emitter 75 μm x 16 cm (New Objective Inc., Woburn, USA), packed in-house with Peprosil C18-AQ resin 3 μm (Dr. Maisch, GmbH, Ammerbuch-Entringen, Germany). LTQ-Orbitrap was set to detect MS spectra at m/z 400 – 2000 with 60,000 resolution at 400 m/z in Orbitrap with a mass lock option m/z 445.120025. Simultaneously, the top 4 most intense ions in MS/MS were acquired in the LTQ with collision induced dissociation. A dynamic exclusion list restricted to 500 m/z values was used for 2 min with a repeat count of 2. Mobile phase A was water/0.1 % FA, mobile phase B 100 % ACN/0.1 % FA. Peptides were separated on linear gradient with mobile phase ramped up to 60 min gradient with 3 to 35 % B was used to elute peptides at constant flow of 300 nl/min.

### LC-MS/MS

For *PRManalysis* - was done using a Q-Exactive (Thermo Fisher Scientific) mass spectrometer connected to an Easy-nLC (Thermo Fisher Scientific). Peptides were first loaded on a trapping column and subsequently separated in-line on a 15 cm C18 column (75 μm ID x 15 cm, ReproSil-Pur 5 μm 200 Å C18-AQ, Dr. Maisch HPLC GmbH, Ammerbuch-Entringen, Germany). PRM scans were acquired at 17500 at 400 m/z resolution with a mass lock option m/z 445.120025, 1e5 AGC target, 27 % collision energy and maximum 50 ms ion accumulation time. 410 ions were monitored in scheduled mode, with each peptide having 3 min analysis window. Mobile phase A was water/0.1 % FA and mobile phase B ACN/water (80:20 (v/v)) with 0.1 % formic acid in a non-linear gradient. B was first increased from 5 to 21 % from 0 to 28 min, then ramped up to 36 % from 28 to 50 min an finally to 100 % from 50 min to 60 min.

### Pre-processing and analysis of AHA LC-MS/MS data

Resulting raw files were searched and quantified with Maxquant 1.6.016 against the UniProt Homo sapiens database (retrieved 12.11.2017) containing isoform information. Carbamidomethylation at cysteine (+57.0215 Da) was set as fixed modification, methionine oxidation (+15.9949 Da) and methionine to AHA substitution (−4.9863 Da) were set as variable modifications for identifications and quantifications. Match between runs and MaxLFQ normalization were turned on. The resulting proteinGroups.txt file was uploaded to Perseus 1.5.8.5, where LFQ intensity values where log2 transformed, and missing values replaced by imputation using Perseus imputation and the normal distribution function. Intensity values were back transformed to original scale, and significantly changing protein groups between LRRK2-G2019S and healthy, and sporadic and healthy were analysed in R. ROTS (An R package for reproducibility-optimized statistical testing, (Elo, et al., 2009)), and Student’s t-test were used with p < 0.05 for significant protein group cut-offs. For both comparisons, significant entries were visualized with a heatmap showing sample intensity values, and all entries were visualized with volcano plots showing p-values and Log2 fold changes.

### Parallel reaction monitoring (PRM)analysis analysis

Raw files from LC-MS/MS analysis were analysed in Skyline. In the output, missing values were replaced by 1 and the data was analysed and visualised using R with the same cut-offs as described for the AHA data analysis with raw intensity values. Statistical analysis was evaluated using Student’s t-test and ROTS (Elo, et al., 2009). Values with p<0.05 and FDR<0.05 were counted as significant and included in the list. For both sporadic vs. healthy and G2019S vs healthy comparisons, significant entries from the list were visualized with a heatmap showing sample intensities, and all entries were visualized with volcano plots showing p-values and Log_2_ fold changes. Enrichment and interaction networks were investigated using the STRING database (Szklarczyk, et al., 2021).

### qPCR analysis

Total RNA from fibroblast cells was extracted using NucleoSpin RNA Mini kit from Macherey-Nagel (#740955) according to the manufacturer’s protocol. The method included DNase I treatment to degrade contaminating genomic DNA. One microgram of total RNA in a 20 μl reaction was reverse transcribed using SensiFast cDNA synthesis kit from Meridian Bioscience (#BIO-65053). Primers for target genes were designed using PrimerQuest™ Tool from Integrated DNA technologies. Quantitative real-time PCR was performed on QuantStudio™ 12K Flex Real-Time PCR System. The reaction mixture (10 μl) contained 1 μl of 30x diluted cDNA (~1.6 ng), 0.5 μl of each primer (10 μM) and 5 μl of PowerUp™ SYBR™ Green Master Mix (# A25742) from ThermoFisher Scientific. Gene expression levels (ΔΔCt) relative to GAPDH were measured and normalized to expression in healthy controls.

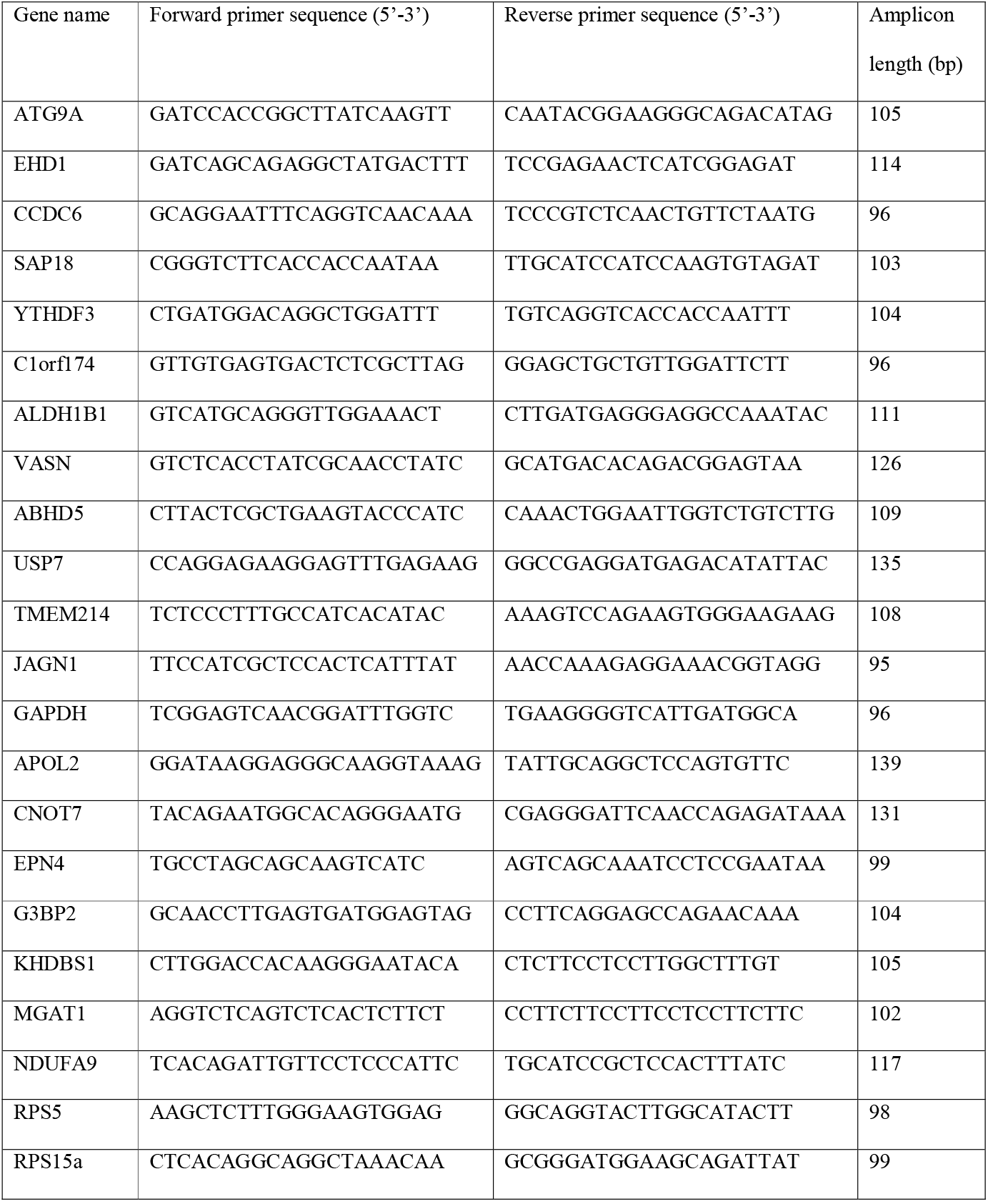

## RESULTS

### Fluorescent non-canonical amino acid tagging (FUNCAT) reveals that nascent protein synthesis is reduced in fibroblasts from sporadic and LRRK2-G2019S Parkinson’s patients

In order to validate our earlier finding that translation is decreased in PD patient cells, we grew non-transformed fibroblasts from patients carrying a LRRK2-G2019S mutation or from individuals diagnosed with sporadic PD, and from healthy donors. Patient details (e.g. age, gender etc.) are described in Table 1. Cells were metabolically labeled using the FUNCAT method as previously described (Deshpande, et al., 2020,tom Dieck, et al., 2015). In this procedure, endogenous methionine was replaced with the azido-hydroxy-homoalanine (AHA) analog. AHA-labeled proteins were then tagged with Alexa-488-alkyne by a cycloaddition *click* reaction (Fig. 1A). This allowed quantitative fluorescence measurement of *de novo* synthesized proteins. Alexa-488 fluorescence was significantly reduced in PD samples compared to control cells indicating that bulk *de novo* synthesis was reduced (Fig. 1B, C). Nascent protein synthesis was reduced in both G2019S carriers and sporadic patients alike.

**Figure 1.**
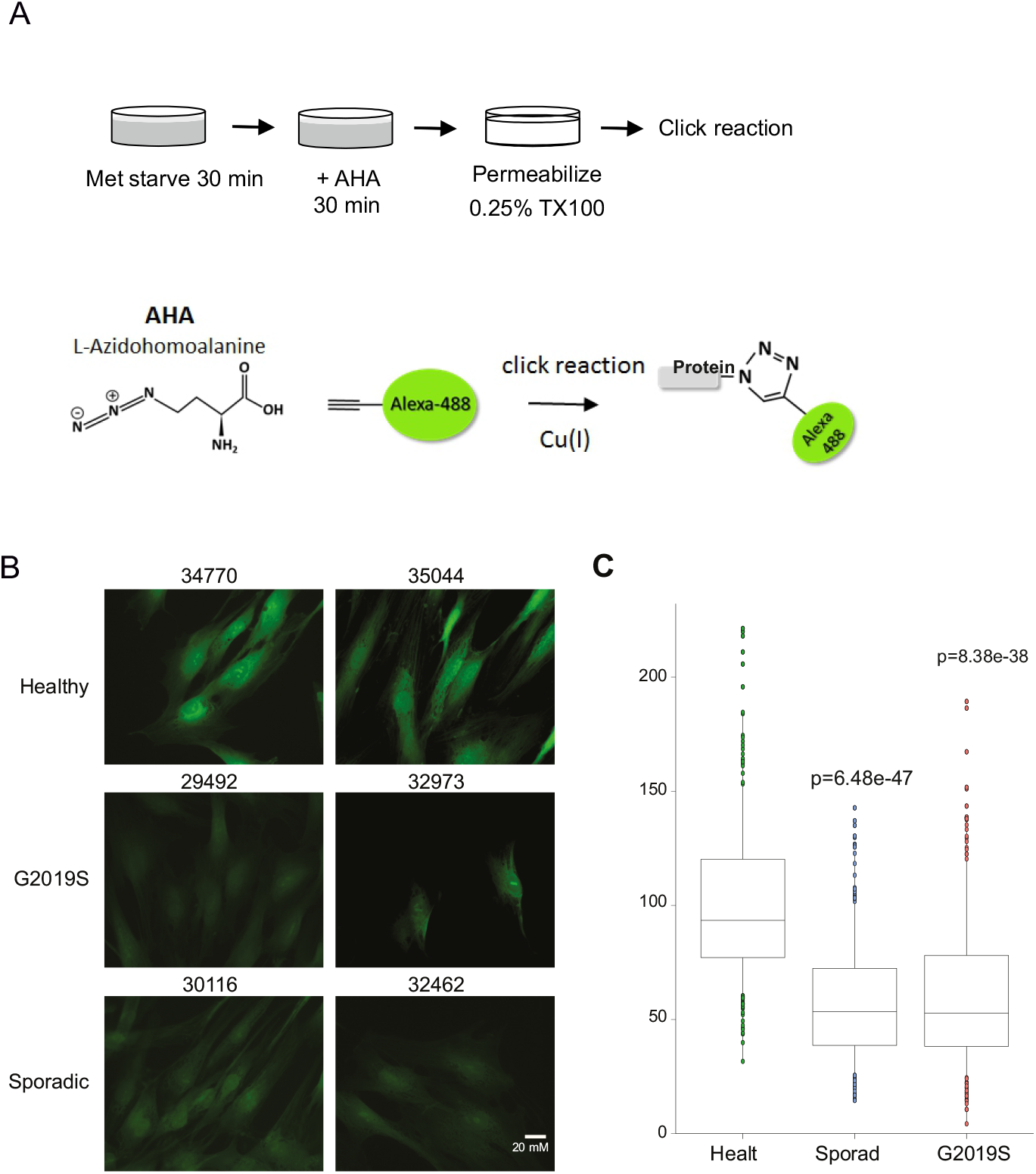
A. Schematic of metabolic labeling. B. Representative images of AHA-labeled fibroblasts cells from G2019S or sporadic Parkinson’s patients or healthy controls and their NINDS repository numbers are shown. C. Quantitative protein synthesis data from healthy, G2019S or sporadic cases are shown. Adjusted p-values were calculated using T-test and corrected with Benjamini and Hochberg method.

### Isolation and LC-MS/MS identification of newly synthesized proteins from LRRK2-G2019S carriers and sporadic patients

We set out to identify which mRNA transcripts were differentially regulated at the level of translation in Parkinson’s patient cells, investigating both sporadic and LRRK2-G2019S cases. Skin punches were isolated from 10 sporadic PD cases and 6 healthy donors from patients attending Turku University Hospital, as previously described (Deshpande, et al., 2020). Five LRRK-G2019S fibroblast samples and 6 healthy controls were from the NINDS and TNGB repositories. Details of subjects including age and sex are described in Table 1.

We used the bio-orthogonal non-canonical amino acid tagging (BONCAT) assay to identify newly translated proteins (Dieterich, et al., 2006) (Fig. 2A). Accordingly, newly synthesized proteins were isolated, and analyzed by mass spectrometry. Data was preprocessed using MaxQuant Perseus analysis software and multiple statistical tests were done using the PhosPiR proteomic and phosphoproteomic data analysis pipeline (Hong, et al., 2021). The sum of peptide intensities for newly synthesized proteins from LRRK2-G2019S and sporadic patients verses healthy are shown in figure 2B and C respectively. The protein intensities were overall lower in cells from PD individuals compared to healthy, regardless of whether they were from LRRK2-G2019S or sporadic patients. Overall, synthesis of 30 nascent proteins were significantly reduced in LRRK2-G2019S cells compared to controls and 33 were significantly decreased in sporadic verses healthy (fig. 2B, C). There was 65% overlap in the identity of proteins that changed significantly between LRRK2-G2019S and sporadic sample groups suggesting a degree of molecular similarity.

**Figure. 2.**
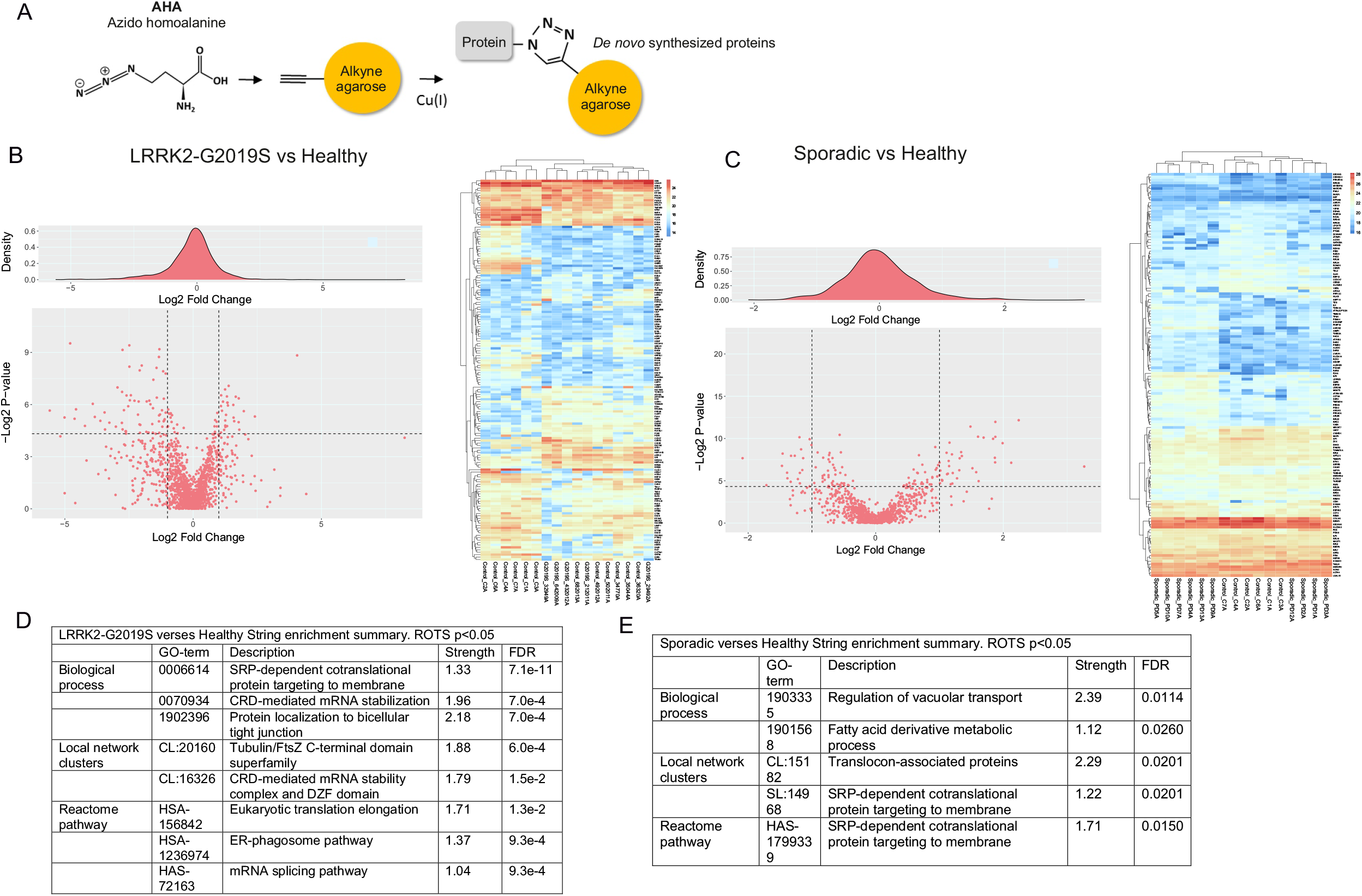
*De novo* protein synthesis is altered in LRRK2-G2019S and sporadic Parkinson’s patient fibroblasts verses healthy controls. A. Schematic overview of click reaction used to isolate newly synthesized proteins using alkyne-linked agarose beads following metabolic labelling with AHA to replace methionine. B. Volcano plot of all AHA-labelled protein intensities for LRKK2-G2019S verses healthy. Heat-map with hierarchical clustering depicts the regulation of nascent protein levels in fibroblasts from LRRK2-G2019S patients verses healthy individuals, union of ROTS and t-test, p-value <0.05. Proteins are labelled with their corresponding gene names in order to save space. C. Volcano plot of all AHA-labelled protein intensities for sporadic verses healthy using ROTS statistical test, and heatmap with hierarchical clustering depicts the nascent proteins as above D, E. Proteins with intensity fold change > 1.5 and p-value < 0.05 were analyzed in STRING. Top ranking functional enrichments are shown for both LRRK2-G2019S and sporadic verses healthy. Strength is calculated from log_10_(observed/expected) network proteins. FDR represents p-values corrected for multiple testing using the Benjamini-Hochberg procedure.

### Signal recognition particle (SRP)-dependent co-translational protein targeting is functionally enriched among nascent proteins that are deregulated in both LRRK2-G2019S and sporadic Parkinson’s

We carried out an enrichment analysis of nascent proteins that were significantly altered in LRRK2-G2019S or sporadic patient cells using the STRING database (Szklarczyk, et al., 2021). This analysis indicated that the biological process “*cytosolic signal recognition particle (SRP)-dependent co-translational protein targeting to membrane*” was functionally enriched among the nascent protein population from LRRK2-G2019S and sporadic patient fibroblasts verses healthy, with FDRs of 7.1E-11 and 1.7E-01 respectively (Fig. 2D, E). This process incorporates components that control the translation of proteins destined for the secretory pathway and their translocation to the endoplasmic reticulum.

In contrast, the Tubulin/FTsz C-terminal domain superfamily network was enriched among nascent proteins from LRRK2-G2019S carriers, but not among sporadic patient nascent proteins (Fig. 2D). This is consistent with the well characterized association of LRRK2 with microtubules (Calogero, et al., 2019).

### Translational targets that were deregulated in Parkinson’s patient cells showed substantially reduced levels in LRRK2-G2019S and sporadic cases alike

We next set out to identify which of the deregulated nascent proteins showed homeostatic disturbance when measured from total cell lysates. For this, we used targeted proteomics and screened for a targeted list of peptides that included unique identifier peptides for all the proteins that were significantly altered in Parkinson’s verses control cells in the BONCAT assay (Fig. 2). The inclusion list incorporated MaxQuant LFQ normalization and data with or without imputation (performed using the Perseus package). Also, two parametric statistical tests were included, Student’s T-test and ROTS. The union list of all significantly changing proteins for G2019S verses healthy and sporadic verses healthy, incorporated 247 proteins in all. The results of the PRM-analysis for healthy verses LRRK2-G2019S groups are shown as volcano plots and heatmaps (Fig. 3).

**Figure 3.**
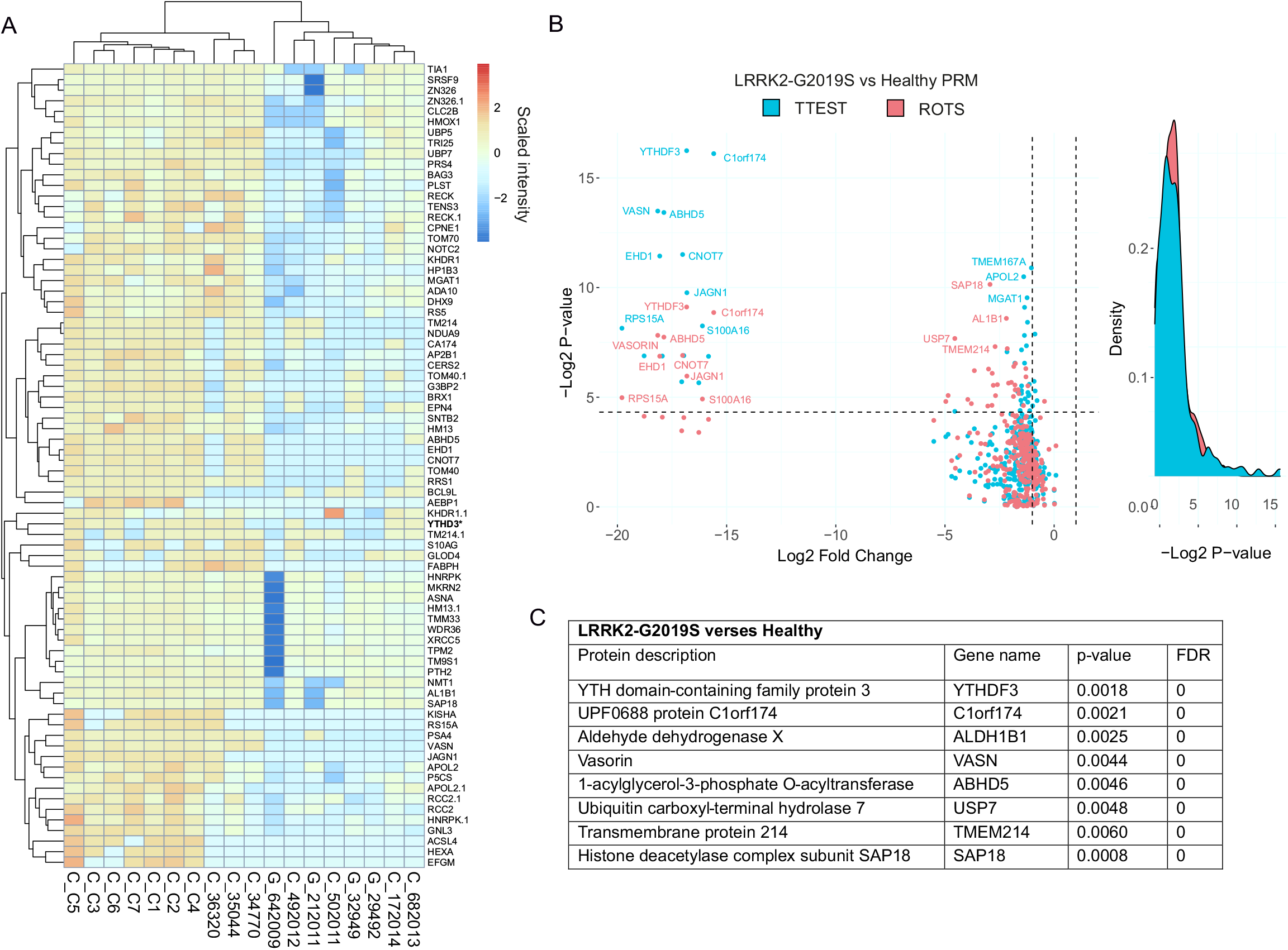
PRM validation of protein homeostasis changes in LRRK2-G2019S patient cells. Targeted proteomics (PRM quantification of peptides from figure 2B) was carried out to validate protein intensity differences observed from AHA-labelled, *de novo* synthesized proteins from total lysates of patient cells. A. Hierarchical clustering of PRM results shows overall protein intensities in fibroblasts from LRRK2-G2019S and healthy controls. Proteins with p-value<0.05 from either ROTS and/or T-test are shown. B. Volcano plot of PRM results from LRRK2-G2019S vs healthy. P-values from ROTS and T-test statistical tests are shown. C. The top changing proteins with FDR=0. from ROTS test are described.

Without exception, all proteins that were significantly changed in the LRRK2-G2019S nascent protein fractions, showed lower levels when measured from total cell lysates using targeted LC-MS/MS validation. Notably, among these proteins, some were substantially decreased by orders of magnitude compared to the levels in fibroblasts from healthy individuals. These proteins were YTH N6-Methyladenosine RNA Binding Protein 3 (YTHDF3), Chromosome 1 open reading frame 174 (C1orf174), Vasorin (VASN), Abhydrolase domain containing 5, lysophosphatidic Acid Acyltransferase (ABHD5), EH domain containing protein 1 (EHD1), CCR-NOT transcription complex subunit 7 (CNOT7), Jagunal homolog 1 (JAGN1), ribosomal protein 15A (RPS15A) and S100 calcium binding protein A16 (S100A16) (Fig. 3B, 3C).

We also performed targeted analysis of whole cell lysates from sporadic patients using the same peptide inclusion list. Sporadic patient samples also showed exclusively decreased intensities compared to those from healthy donors (Fig. 4A-C), although the size effect was lower than in the LRRK2-G2019S patient group.

**Figure 4.**
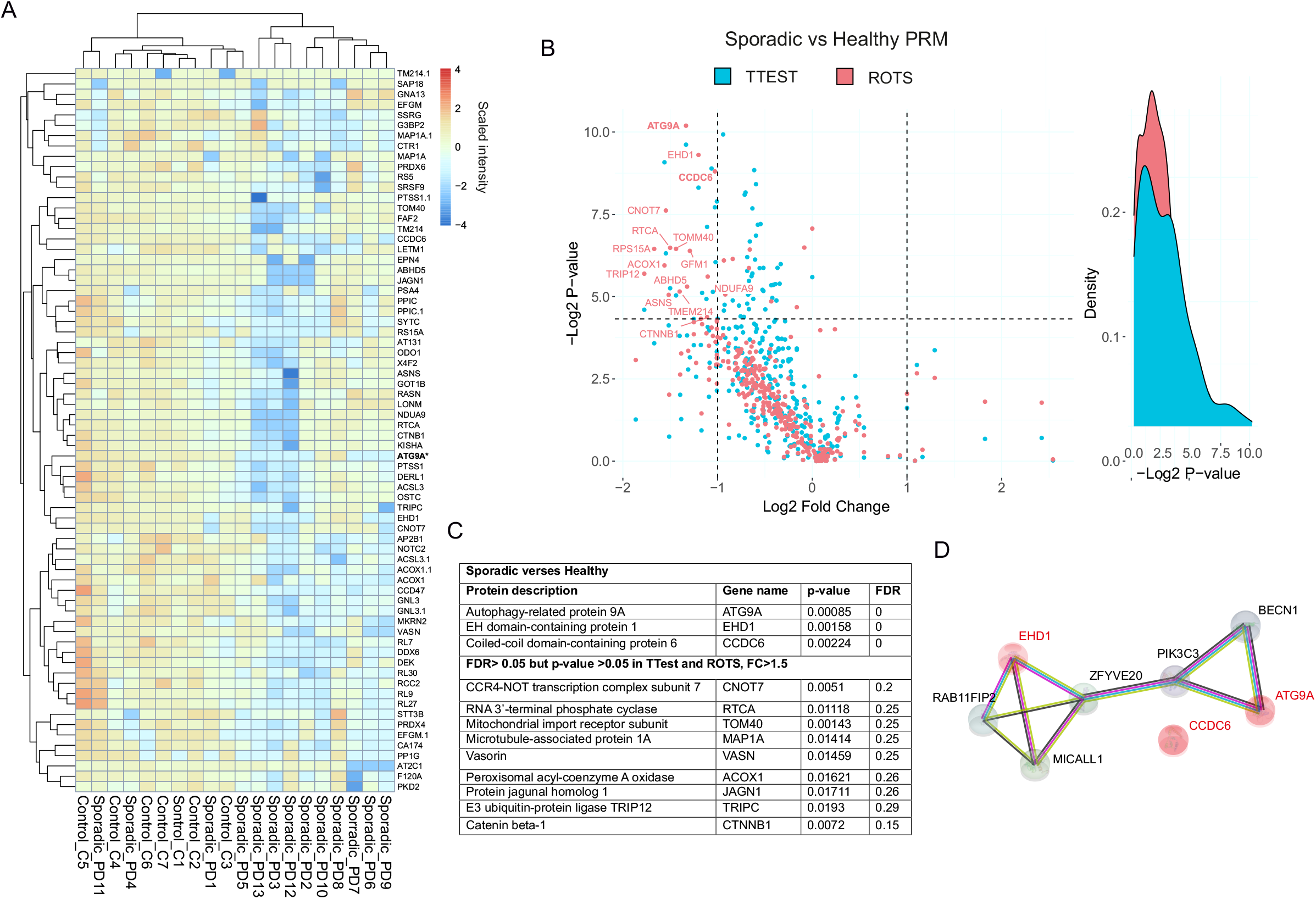
PRM validation of protein homeostasis changes in sporadic Parkinson’s patient cells. Targeted proteomics (PRM quantification of peptides from figure 2C), was carried out to validate protein intensity differences from total lysates of patient cells. A. Hierarchical clustering of PRM results shows overall protein intensities in fibroblasts from sporadic and healthy controls with a p-value <0.05 in either ROTS and/or T-test. B. Volcano plot of PRM results from sporadic vs healthy. P-values from ROTS and T-test statistical tests are shown. C. The top changing proteins are described. D. STRING network showing connectivity of ATG9A and EHD1.

### Several of the proteins that were decreased in LRRK2-G2019S patient cells were also decreased in cells from sporadic patients

The question as to whether LRRK2-G2019S Parkinson’s and sporadic Parkinson’s are molecularly overlapping or whether they represent molecularly distinct forms of disease is of interest (Deshpande, et al., 2020,Di Maio, et al., 2018,Melachroinou, et al., 2020). We therefore examined the overlap in proteins that showed altered levels from the LRRK2-G2019S and sporadic patient groups. The overlap between these Parkinson’s subtypes is shown in the Venn diagram in figure 5A. In the sporadic Parkinson’s group, almost all deregulated proteins overlapped with the LRRK2-G2019S group with the exception of the autophagy-related protein 9A (ATG9A) and CCDC6 (Fig. 5A). ATG9A plays a key role in autophagy, specifically in the formation of the pre-autophagosomal structure assembly site and in vacuole transport vesicle formation (Puri, et al., 2014). In LRRK2-G2019S patients, the protein that was most highly downregulated was YTHDF3, which was unique to this patient group. This protein belongs to a family of N6-methyladenosine RNA modification readers that can recruit molecular complexes to M^6^A sites, to impact RNA splicing, export, stability, trafficking and translation efficiency (Yen and Chen, 2021). Functional enrichment for the LRRK2-G2019S patient data identified mRNA splicing (GO terms 0048024, 0043484 and CL16101), but also pre-ribosome and ribosome biogenesis (GO term CL14573) as the most highly enriched functions. In contrast, in the sporadic Parkinson’s group, SRP-dependent and viral mRNA translation processes (GO term 0006614) and viral mRNA translation and peptide chain elongation as well as the ribosome KEGG pathway (GO terms CL14985, CL14976 and HSA03010) were enriched (Fig. 5B).

**Figure 5.**
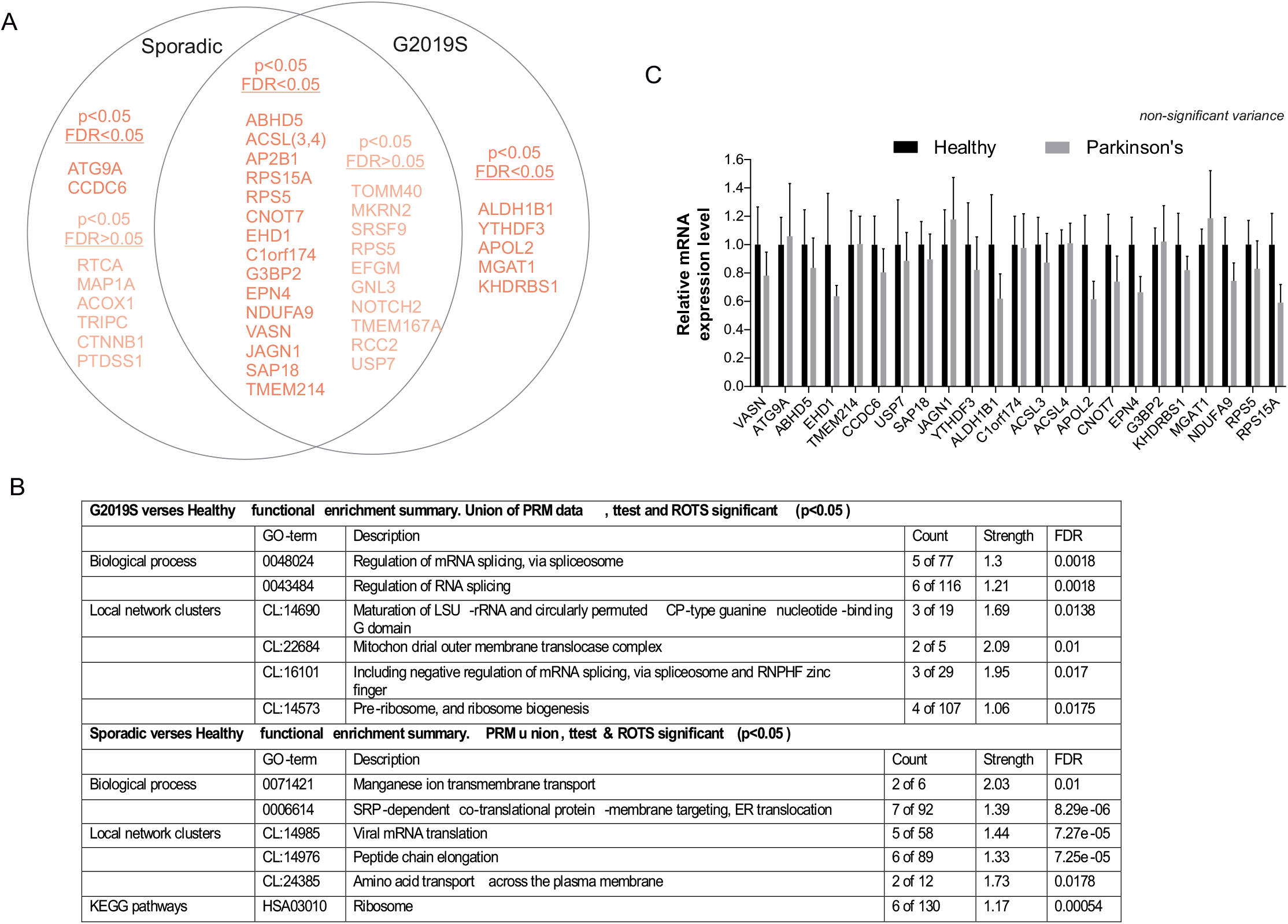
A. Venn diagram summary of proteins that show reduced homeostasis in LRRK2-G2019S and sporadic patients. Proteins (indicated by gene names) are grouped according to significance. For each group comparison, proteins that have P<0.05 and FDR<0.05 in either ROTS or T-test are bolded. Proteins with p<0.05 but FDR>0.05 in either test are shown below in a lighter shade. Proteins belonging to the overlapping group are those with FDR<0.05 in either ROTS or T-test for sporadic or LRRK2-G2019S and FDR<0.05 or p<0.05 in the other Parkinson’s type. B. Q-PCR was carried out to determine if disturbed homeostasis was due to reduced mRNA levels in sporadic patient cells (n=13) verses healthy (n=7), rather than disrupted translation. Mean data normalized to gene expression in healthy +/- S.E.M. is shown. Student’s t-test did not reveal significant changes in the gene expression. C. STRING enrichment analysis of significantly changing proteins in LRRK2-G2019S and sporadic Parkinson’s disease. Top ranking functional enrichments for biological process, local network clusters and KEGG pathways are shown. Input data is from the union of ROTS and ttest PRM analysis changes (p<0.05). Strength is calculated from log_10_(observed/expected) network proteins. FDR values show p-values corrected for multiple testing using the Benjamini-Hochberg procedure.

In case any of the protein expression changes in patient cells were due to reduced mRNA levels, we carried out quantitative PCR for the mRNAs of the most significantly changing proteins. This analysis failed to detect any significant changes in mRNA levels (Fig. 5C). These data indicate that there is a deficit in the synthesis of certain proteins in fibroblasts obtained from LRRK2-G2019S and sporadic Parkinson’s patients relative to healthy controls. This results in overall reduced levels in a subset of proteins, many of which are reduced in both patient groups.

## DISCUSSION

Dysregulation of protein synthesis has been associated with Parkinson’s disease through a variety of studies in patient-derived cells and microphysiological systems (Deshpande, et al., 2020,Kim, et al., 2021,Trapecar, et al., 2021). Here, we set out to identify which proteins were deregulated at the level of translation in patient cells using the BONCAT method (Dieterich, et al., 2006). This method uses proteomics to identify nascent proteins that have been purified based on a non-canonical amino acid tag that is introduced to the cells during a pulse labeling. We examined nascent proteins from fibroblasts isolated from the sporadic Parkinson’s cases and healthy volunteers from Southwest Finland, and from cells of patients carrying a gain of function LRRK2-G2019S mutation, obtained from the NINDS and TNGB repositories. Discovery phase analysis identified that several of the nascent proteins differed in intensity in patients cells. Functional enrichment analysis of these differentially expressed nascent proteins identified that SRP-targeted translation was enriched in both sporadic and LRRK2-G2019S patient groups. Interestingly, Tubulin /FtsZ pathway (CL:20160) was also enriched in the LRRK2-G2019S group as was the endoplasmic reticulum-phagosome pathway (HSA-1236974), consistent with the implicated deficits in these cellular processes in Parkinson’s pathology (Bonam, et al., 2021,Watanabe, et al., 2020).

Altered protein synthesis *per se* will not necessarily manifest in altered protein expression let alone functional deficit. The cell is equipped with proteostatic mechanisms that can compensate for deficiencies in synthesis by controlling turnover for example. It was therefore important for us to establish whether any of the regulated nascent proteins were altered at the whole cell level. Targeted proteomic analysis revealed that a subset of proteins were significantly decreased at the level of the whole fibroblast proteome and a number of these changes occurred both in sporadic and LRRK2-G2019S patients. Of note, the mRNA levels for these proteins were unchanged, consistent with a deficit in protein synthesis playing a causal role. It is worth noting that the data here may underestimate the truly changing proteins as even targeted proteomic analysis can suffer from false negatives due to undersampling (Schmidt, et al., 2008). Also, the Parkinson’s patients lysates showed consistently lower overall protein concentrations which were normalized prior to mass spectrometry analysis, providing potential for underestimation of decreases. Nonetheless, we focus here on the proteins that remained significantly changed, some substantially so, in patient fibroblast proteomes. Many of these are functionally interesting in the context of Parkinson’s pathology, as discussed below.

We find that the top downregulated proteins in sporadic patient fibroblasts were autophagy-related protein-9A (ATG9A) and EHD1, which is also involved in retromer formation and endosomal sorting (Cui, et al., 2019), and CCDC6, a proposed tumor suppressor that was recently implicated as a risk gene for Alzheimer’s disease (Schwartzentruber, et al., 2021). Autophagy is the process whereby cells target cellular waste, such as aggregated or misfolded proteins for degradation by the lysosome. A deficiency in autophagy is believed to contribute to the core component of Parkinson’s pathology, i.e. the formation of alpha-synuclein aggregates (Xilouri, et al., 2016). We show that the mRNA for these proteins is unchanged in patient fibroblasts compared to controls, yet protein levels are reduced both when measured at the level of nascent proteins in the BONCAT assay, and when measured at the level of whole cell proteome. This suggests that there may be a deficit in the synthesis of these proteins in sporadic patient cells. ATG9A is an essential player in autophagy and mitophagy, being required for autophagosome assembly and maturation, where it directs the supply of membrane to maturing autophagosomes (De Tito, et al., 2020,Levine and Kroemer, 2019,Yamano and Youle, 2020). Interestingly, ATG9A, is also implicated in the innate immune response (Saitoh, et al., 2009). We observed a 2.5 fold decrease in levels of ATG9A on average, but when one examines individual samples, some patients have undetectable levels. Interesting, in LRRK2-G2019S samples, we did not detect a significant decrease in ATG9A, even though LRRK2-G2019S, among other Parkinson’s genes glucocerebrosidase and PINK1, encode mutations that lead to reduced autophagy (Bonam, et al., 2021,Levine and Kroemer, 2019). Moreover, in fibroblasts from LRRK2-G2019S and other familial Parkinson’s cases, autophagic flux was shown to be reduced (González-Casacuberta, et al., 2019, Korecka, et al.). Therefore, we were surprised that ATG9A and EHD1 were not significantly altered in LRRK2-G2019S patients. However, it is worth noting that there were only four patients available for LRRK2-G2019S and ATG9A was reduced in 75% of cases, but this was not sufficient to give statistical significance. Thus, it is possible that ATG9A is also a relevant target for downregulation in LRRK2-G2019S. This is the first report to our knowledge of ATG9A dysregulation in Parkinson’s. It would be interesting to find out if it is downregulated in other tissues, for example in the gut or brain, as this could explain the build-up of alpha-synuclein aggregates. It would however be important for independent studies to replicate these findings.

In the LRRK2-G2019S cohort, YTHDF3 was the most strongly downregulated protein, whereas in sporadic cases, YTHDF3 was decreased in 44% of samples. YTHDF3 recognizes the common M6-Methyladenosine modification of mRNA which acts to promote mRNA splicing, its stability and translation efficiency (Wang, et al., 2015). Down regulation of YTHDF3 is expected to reduce translation. Consistent with this, enrichment analysis identified biological processes involving RNA splicing (FDR 0.0018) and large subunit ribosomal RNA maturation (FDR= 0.0138) as a general theme among the stably downregulated proteins. On the other hand, in sporadic patients, functional enrichment identified manganese ion transport (FDR=0.01) and various types of translation regulatory processes (FDR=8.29 E-06), including peptide chain elongation (FDR=7.25 E-05) and viral mRNA translation (FDR=7.25 E-05). Thus there was a predicted functional overlap at the level of protein translation in both disease subgroups. This was born out by the considerable overlap in downregulated proteins in sporadic and LRRK2-G019S carriers.

In summary, analysis of Parkinson’s patient fibroblasts identified key regulators of autophagy, translation and endosomal sorting as downregulated at the stage of protein synthesis. Our analysis identifies molecular commonality between sporadic and LRRK2-G2019S patients at the level of function, and some of the specific downregulated targets are shared in both subtypes. Of specific interest are the proteins that change with the highest level of statistical significance, ATG9A, a requisite autophagy protein, and YTHDF3, a regulator of M6 adenosine methylation. Although this study is carried out in patient skin cells, the changes observed may be relevant for brain, which like fibroblasts derives originally from ectoderm and retains potential to differentiate to neuronal cell types. Furthermore, these results could provide useful PD biomarkers, especially if the less invasive skin scrape biopsies were to come into use.

## ACKNOWLEDGEMENTS

Susanna Pyökäri is acknowledged for assisting with fibroblast cultures and Veronica Fagerholm for sample management. This work was funded by Business Finland project # 1817/31/2015 (PD, PJ, DF, and YH), Åbo Akademi University (E.C), and the Michael J. Fox Foundation (DF, YH) and the MATTI graduate school (YH). We acknowledge the facilities and staff of the Cell Imaging Cytometry core (CIC) and Turku Proteomics Facility which are supported by Biocenter Finland.

## Notes

### Competing Interest Statement

The authors have declared no competing interest.

